# Hierarchical classification of hematologic malignancies using epigenetic and genetic information

**DOI:** 10.64898/2026.07.02.735835

**Authors:** Maximilian Schönung, Melissa Türe, Panna Lajer, Simon Renders, Tobias Rausch, Til L. Steinicke, Anna Dolnik, Eric Sträng, Meghana S. Oak, Jessica Heilmann, Kevin Roth, Lena Katzenstein, Christian Rohde, Etienne Sollier, Peter Horak, Tim Sauer, Jonathan C. Strefford, Martí Duran-Ferrer, Christopher C. Oakes, José Ignacio Martín-Subero, Ulrich Germing, Michael Dworzak, Albert Catala, Christian Flotho, Charlotte M. Niemeyer, Hartmut Döhner, Volker Hovestadt, Stefan Fröhling, Richard F. Schlenk, Florian H. Heidel, Jan Korbel, Clarissa Gerhäuser, Mark Hartmann, Carsten Müller-Tidow, Pavlo Lutsik, Michael Hundemer, Miriam Erlacher, Lars Bullinger, Christoph Plass, Daniel B. Lipka

## Abstract

Molecular testing in hematology requires different assays for disease subgroup identification, risk stratification and selection of appropriate treatment regimens. Yet, molecular tests are not necessarily standardized between diagnostic laboratories, resulting in varying turnaround times and potentially divergent results. To resolve this issue and enable single-assay molecular testing, we have developed a hierarchical classification framework that combines epigenetic and genetic data from whole genome nanopore sequencing (WGNS) with machine learning to determine disease entities, epigenetic subgroups (epitypes) and genetic aberrations in hematopoietic neoplasms. We curated DNA methylation data from 5,420 samples and trained a classifier allowing entity-level diagnostics featuring 21 conditions, including healthy controls, acute and chronic myeloid and lymphoid neoplasms. This classifier was subsequently combined with entity-specific epitype classifiers predicting 44 therapeutically or prognostically relevant states, followed by integration of genetic data. Benchmarking of the combined (epi-)genetic testing strategy using WGNS confirmed high accuracy in the detection of diagnostic groups and risk stratification, and identified diagnosis-defining molecular alterations that were not reported by standard-of-care work-up.

## INTRODUCTION

Over the past two decades, molecular diagnostics have enabled improved risk classification and identification of targetable alterations in hematopoietic neoplasms^1–5^. These discoveries have largely been based on genetic and transcriptomic data, due to the widespread availability of these technologies across institutes, the development of standardized workflows, and the capacity to detect genetic driver events. However, while genetic driver events can be shared across different disease entities, the DNA methylome intrinsically encodes cell-, lineage-, and disease-specific patterns, thus potentially enabling a more accurate identification of molecular disease subgroups. In line with this, DNA methylation analysis has recently been adopted for the molecular diagnosis of brain cancers and sarcomas, and is now being used increasingly in molecular tumor boards for solid cancers^6–9^. In hematopoietic neoplasms, DNA methylation analysis has been used to identify molecular subgroups in various disease entities such as acute lymphoblastic leukemia (ALL), acute myeloid leukemia (AML), chronic lymphocytic leukemia (CLL), chronic myelomonocytic leukemia (CMML), myelodysplastic syndrome (MDS), splenic marginal zone lymphoma (SMZL), and juvenile myelomonocytic leukemia (JMML)^10–20^. However, most DNA methylation-based disease subgroups have not been translated into clinically applicable diagnostic tools or workflows. We recently sought to address this limitation for JMML by establishing an international consensus on the definition of DNA methylation subgroups, so-called epitypes, in this disease, and by developing clinically applicable tools that enable the faithful classification of individual patients^21^. This effort has resulted in the first clinical trial that stratifies JMML patients by epitype to receive different treatments according to their predicted risk (NCT05849662)^22^.

More recently, two DNA methylation classifiers spanning different types of acute leukemia have been developed to identify molecular subgroups, which can be used in combination with nanopore sequencing^23,24^. These classifiers enable rapid molecular diagnostics with improved turnaround times compared to the current standard-of-care. However, these classifiers only allow valid interpretation of prediction results for patients for which a diagnosis of acute leukemia has been established. Currently, this requires additional laboratory tests to first establish an entity-level diagnosis, followed by the selection of an appropriate classifier model. Here, we aimed to overcome this limitation by enabling combined entity and disease subgroup classification from DNA methylation data by using a hierarchical classification strategy. First, this strategy establishes the diagnosis of a disease entity using a broad DNA methylation classifier comprising 21 different entities and controls. Based on the prediction result, we then select an appropriate downstream classifier for identification of a molecular subgroup. Combining this classification strategy with multi-omics whole genome nanopore sequencing (WGNS) enables further genetic testing and provides a complete set of molecular data necessary for clinical decision making.

## RESULTS

### DNA methylation reference atlas of hematopoietic neoplasms and healthy cell types

We collected DNA methylation array data of adult and pediatric hematopoietic neoplasms and healthy controls, comprising a total of 7,053 samples (**Figure 1A**). A stringent quality control pipeline was applied to remove probes mapping to sex chromosomes, non-CpG targets, common single nucleotide polymorphisms (SNPs), as well as previously annotated poor quality probes and probes with low detection p-values^25^. Samples with low-quality methylation array data and duplicated samples that were included in multiple published datasets were removed, resulting in a final dataset of 5,420 samples (3,796 hematopoietic neoplasms and 1,624 controls; **Supp. Table 1&2**). This comprised 20 different disease entities, encompassing myeloid and lymphoid acute and chronic neoplasms (i.e. acute lymphoblastic leukemia (ALL), acute myeloid leukemia (AML), mixed-phenotype acute leukemia (MPAL), chronic myeloid leukemia (CML), chronic myelomonocytic leukemia (CMML), juvenile myelomonocytic leukemia (JMML), myelodysplastic syndrome (MDS), myelofibrosis (MF), essential thrombocythemia (ET), polycythemia vera (PV), Noonan-syndrome associated myeloproliferative disorder (NS-MPD), Burkitt lymphoma (BL), diffuse large B-cell lymphoma (DLBCL), follicular lymphoma (FL), mantle cell lymphoma (MCL), splenic marginal zone lymphoma (SMZL), chronic lymphocytic leukemia (CLL), B-cell prolymphocytic leukemia (BPLL) and Waldenström macroglobulinemia (WM) patients). Samples of isolated hematopoietic cell types from blood, bone marrow (BM), tonsils and spleen (n=578), as well as healthy blood and cord blood samples (n=1,046) were used as controls (**Figure 1B**). Unsupervised dimension reduction was performed based on variable CpG sites (n=12,627 varCpGs; **Supp. Table 3**).

**Figure 1.**
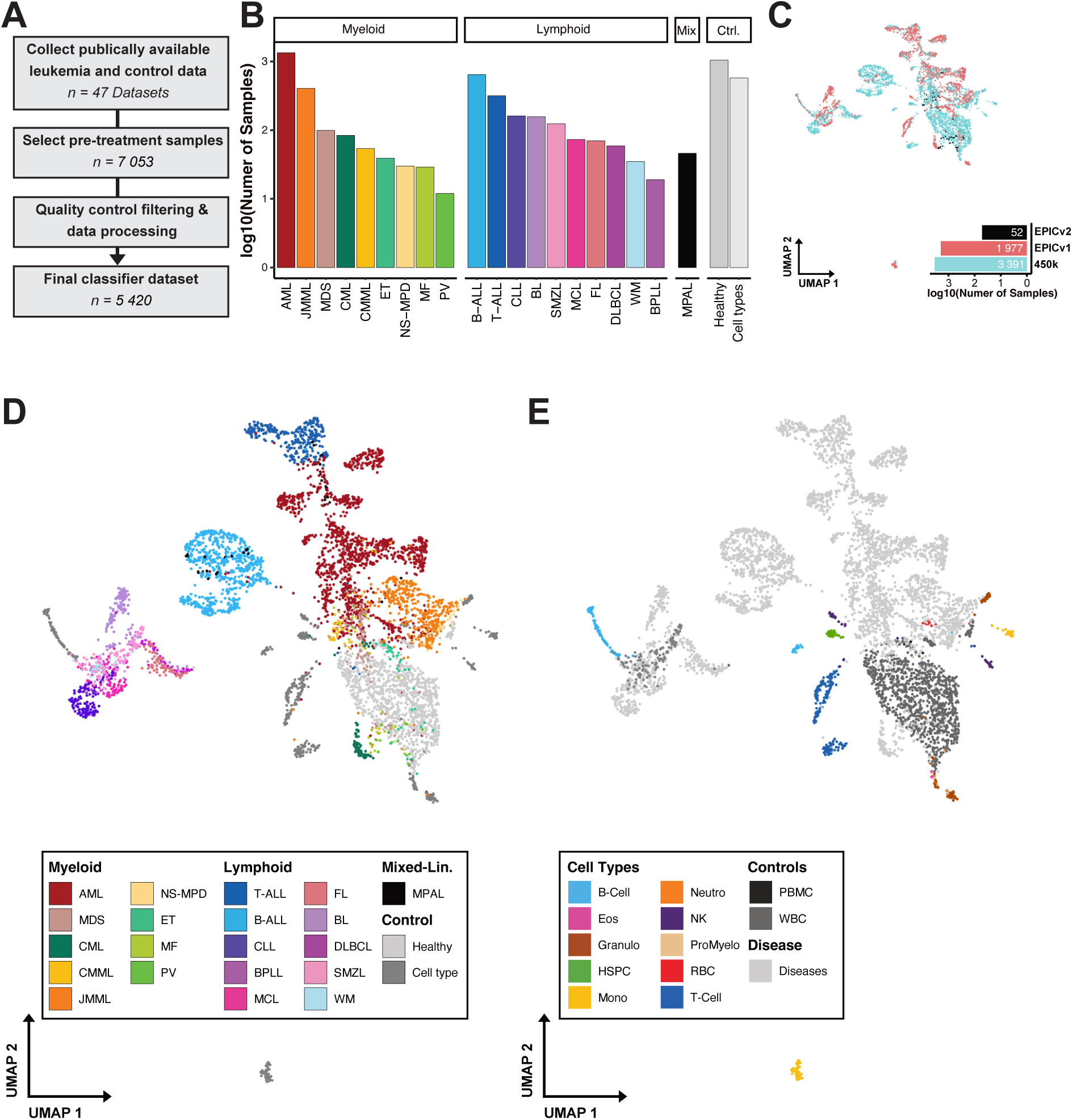
Global DNA methylation patterns separate hematopoietic neoplasms from healthy cell types. **A)** Overview of the sample processing workflow. **B)** Number of included samples after quality control and processing. Barplot showing the log10 sample number per entity. **(C-E)** UMAP dimension reduction analysis using variable CpG sites (n= 12,627) colored by DNA methylation array technology (**C**), disease entities **(D)** and healthy cell types **(E)**. (AML, acute myeloid leukemia; B-ALL, B-cell acute lymphoblastic leukemia; BL, Burkitt lymphoma; BPLL, B-cell prolymphocytic leukemia; CLL, chronic lymphocytic leukemia; CMML, chronic myelomonocytic leukemia; CML, chronic myeloid leukemia; DLBCL, diffuse large B-cell lymphoma; Eos, eosinophils; ET, essential thrombocythemia; FL, follicular lymphoma; Granulo, granulocytes; Healthy, healthy control; HSPC, hematopoietic stem and progenitor cells; JMML, juvenile myelomonocytic leukemia; MCL, mantle cell lymphoma; MDS, myelodysplastic syndrome; MF, myelofibrosis; Mono, monocytes; MPAL, mixed-phenotype acute leukemia; Neutro, neutrophils; NK, natural killer cells; NS-MPD, Noonan syndrome-related myeloproliferative disorder; PBMC, peripheral blood mononuclear cells; ProMyelo, promyelocytes; PV, polycythemia vera; RBC, red blood cells; SMZL, splenic marginal zone lymphoma; T-ALL, T-cell acute lymphoblastic leukemia; WM, Waldenström macroglobulinemia; WBC, white blood cells)

Annotation of disease entity and cell type identity in the reduced dimensionality space revealed entity- and cell type-specific clustering of samples, with no evident batch effects caused by differences in DNA methylation array technology (**Figure 1C-E**). To quantify the degree of differentiation versus tumor-specific DNA methylation changes in these samples for a comparative analysis, we identified differentially methylated probes (DMPs) between disease entities (entity DMPs, n= 135,888) and between healthy cell types (cell type DMPs, n=32,744). DMP were subsequently categorized as shared (n=26,493; 19%), differentiation- (n=6,251; 4%) and tumor-specific (n=109,395; 77%) based on the overlap between the two sets (**Supp. Figure 1A; Supp. Table 4**). To better understand the biological programs that are differentially regulated by DNA methylation during either physiological differentiation or transformation, we analyzed ChromHMM chromatin state enrichment in each DMP subset (**Supp. Figure 1B**). We observed an enrichment of active enhancers in cell type DMPs and in shared DMPs that overlap between cell type and entity DMPs. In contrast, tumor-specific DMPs were enriched for bivalent transcription start sites, bivalent enhancers and polycomb-repressed regions.

Thus, our data support a model in which the DNA methylome of hematopoietic neoplasms reflects a combination of differentiation-specific patterns, reflecting cellular identity, which is programmed at enhancers, and additional tumor-specific patterns, which mainly involve bivalent chromatin. The unsupervised analysis of DNA methylation patterns indicated that these patterns can discriminate hematologic neoplasms and potentially qualify as epigenetic signatures that can be used to train a classifier to diagnose such diseases.

### Development of a DNA methylation classifier for hematopoietic neoplasms

In a next step, we aimed to train a DNA methylation classifier that enables prediction of hematopoietic disease entities based on our reference atlas. After excluding isolated cell types, the dataset for classifier training consisted of 4,842 samples, comprising 21 classes (20 hematopoietic neoplasms and healthy controls). The data was divided into a training (n=3,884) and testing (n=958) cohort while ensuring balanced class distributions (**Figure 2A**). An extreme gradient boosting (XGBoost) model was trained based on the previously determined varCpGs (n=12,627), followed by hyperparameter tuning and feature selection. The final classifier comprised 932 CpGs and demonstrated an entity classification accuracy of 95.5% with a median class specificity of 1 (min: 0.98; max: 1), and sensitivity of 0.93 (min: 0; max: 1; **Supp. Figure 2A&B; Supp. Table 5**). The per sample class (entity) prediction probabilities ranged from 0.21 to 1.0 (mean: 0.96; median: 1). To determine the prediction probability threshold required to accurately assign patients to each class, we calculated a binarized ROC curve based on correctly and incorrectly assigned patients using the maximum class accuracy (**Figure 2B**). This analysis revealed that a class probability threshold of 0.972 resulted in the highest classifier sensitivity and specificity (**Figure 2B, Supp. Figure 2C&D**). Using this cut-off, the classifier achieved prediction accuracy of 99.4% while labeling 20% (n=103) of patients who did not meet the probability threshold as “unknown”. We used this cut-off as a “tier-1” filter, in which the model predicts the disease entities with the highest prediction accuracy.

**Figure 2.**
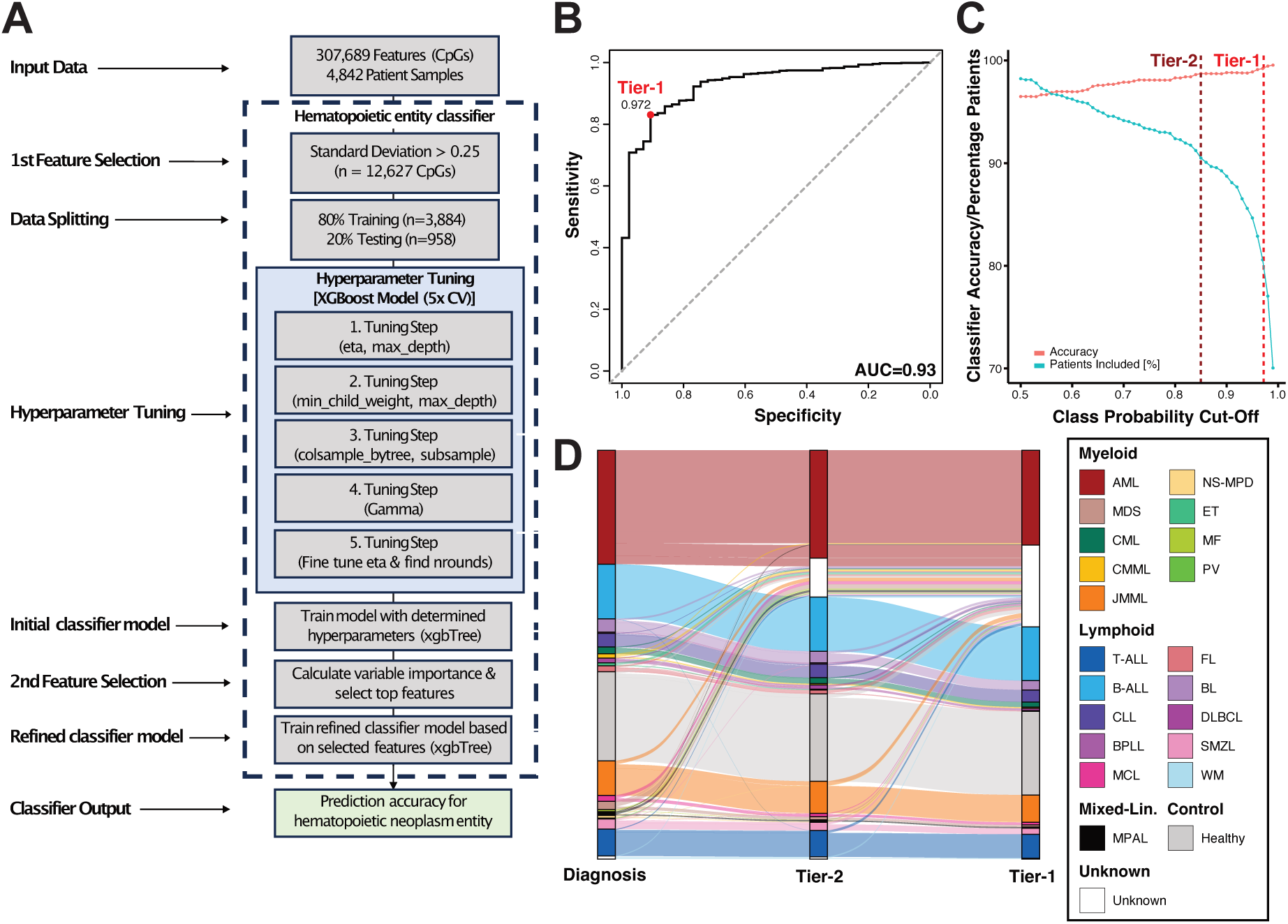
Development of a DNA methylation classifier for the diagnosis of hematopoietic neoplasms. **(A)** Schematic overview summarizing the different steps of classifier development. **(B)** ROC curve based on maximum class probability for binarized predictions. Area under the curve (AUC) and best performance threshold (Tier-1) are annotated. **(C)** Classifier accuracy (red dots and line) and respective percentage of patients with valid predictions (prediction probability above each cut-off; blue dots and line) are plotted. Tier-1 and tier-2 accuracy cut-offs are annotated as dashed lines. **(D)** Alluvial plot comparing the original diagnosis for the testing cohort (n = 958 patients) to the classification results. Bars represent the fraction of patients with the respective diagnosis and connecting lines the concordance between original diagnosis and classifier predictions. Two different prediction probability cut-off levels (tier-1 and tier-2) are used and patients that do not reach this cut-off are assigned to “unknown”.

However, as this stringent cut-off substantially reduced the number of patients with successful entity predictions, we additionally introduced a “tier-2” probability threshold of 0.85, which was selected based on a trade-off analysis between number of included patients and overall classifier accuracy (**Figure 2C**). The “tier-2” filter results in a classifier accuracy of 98.7% and allows us to predict 91% of patients (**Figure 2C-D, Supp. Figure 2E&F**). Based on these cutoffs, the model predicted the 21 entity classes with high specificity (tier-1: median 1, min. 0.99, max. 1; tier-2: median 1, min. 0.99, max. 1) and sensitivity (tier-1: median 1, min. 0, max. 1; tier-2: median 1, min. 0.5, max. 1) across both confidence tiers, thus providing a framework for accurate DNA methylation-based entity classification for hematologic malignancies.

### Hierarchical epigenetic classification of hematopoietic neoplasm disease entity and epitype

Previously published DNA methylation classifiers required prior laboratory testing to first determine an entity-level diagnosis before selecting an appropriate epigenetic subgroup (epitype) classifier. Our newly established classifier addresses this limitation by directly inferring the respective disease entity for hematopoietic neoplasms from DNA methylation data. This enables a hierarchical classification strategy in which first the respective disease entity is predicted, and then, based on the prediction results, an appropriate epitype classifier is determined and applied to establish a DNA methylation-based entity and epitype diagnosis (**Figure 3A**). We validated this strategy for B-ALL, CLL and JMML, as DNA methylation epitypes have been well-established for these diseases. Validation was conducted in cohorts which had not been included in classifier training, allowing an unbiased assessment of our classifier in a real-world scenario. For B-ALL, we used a publicly available dataset^26^ and when applying the tier-1 cut-off, 92 out of 133 patients (69.2%) were classified as B-ALL, while 41 out of 133 patients (30.8%) did not reach the prediction cutoff (**Figure 3B; Supp. Table 6**). Reanalyzing these patients using the tier-2 threshold, correctly classified 114/133 patients (85.7%) as B-ALL and 19/133 (14.3%) as unknown. Epitypes were subsequently determined for all patients that were classified as B-ALL according to the tier-2 threshold. We therefore relied on MARLIN, a recently developed epitype model for acute leukemias^23^. However, as the originally published neural network-based MARLIN model was developed for sparse genome-wide sequencing data, we developed a random-forest based MARLIN model (MARLIN-RF) for improved prediction accuracy with non-sparse array-based data. Five-fold cross validation accuracies for MARLIN methylation classes based on this model were 0.96 for AML, 0.95 for B-ALL, 0.94 for T-ALL, 0.95 for MPAL and 0.99 for controls. Using this MARLIN-RF classifier, 57 B-ALL samples (50%) were classified as high hyperploid (HeH), 44 (38.6%) as *ETV6*::*RUNX1*, 3 (2.6%) as *TCF3*::*PBX1/MEF2D*-r and 1 (0.9%) patient as *KMT2A*-rearranged epitypes (**Figure 3B**). Nine patients (7.9%) did not reach the MARLIN cutoff of 0.8 and were classified as unknown. Considering the published genetic groups for the patients in the B-ALL dataset, all 57 (100%) HeH epitype patients were annotated as hyperdiploid, and 43/44 (97.7%) *ETV6*::*RUNX1*, 1/1 (100%) *KMT2A*-rearranged and 2/3 (66.7%) *TCF3*::*PBX1/MEF2D*-r epitype assigned patients had the respective defining genetic alterations, providing genetic support for the respective B-ALL epitype assignments (**Supp. Figure 3A**).

**Figure 3.**
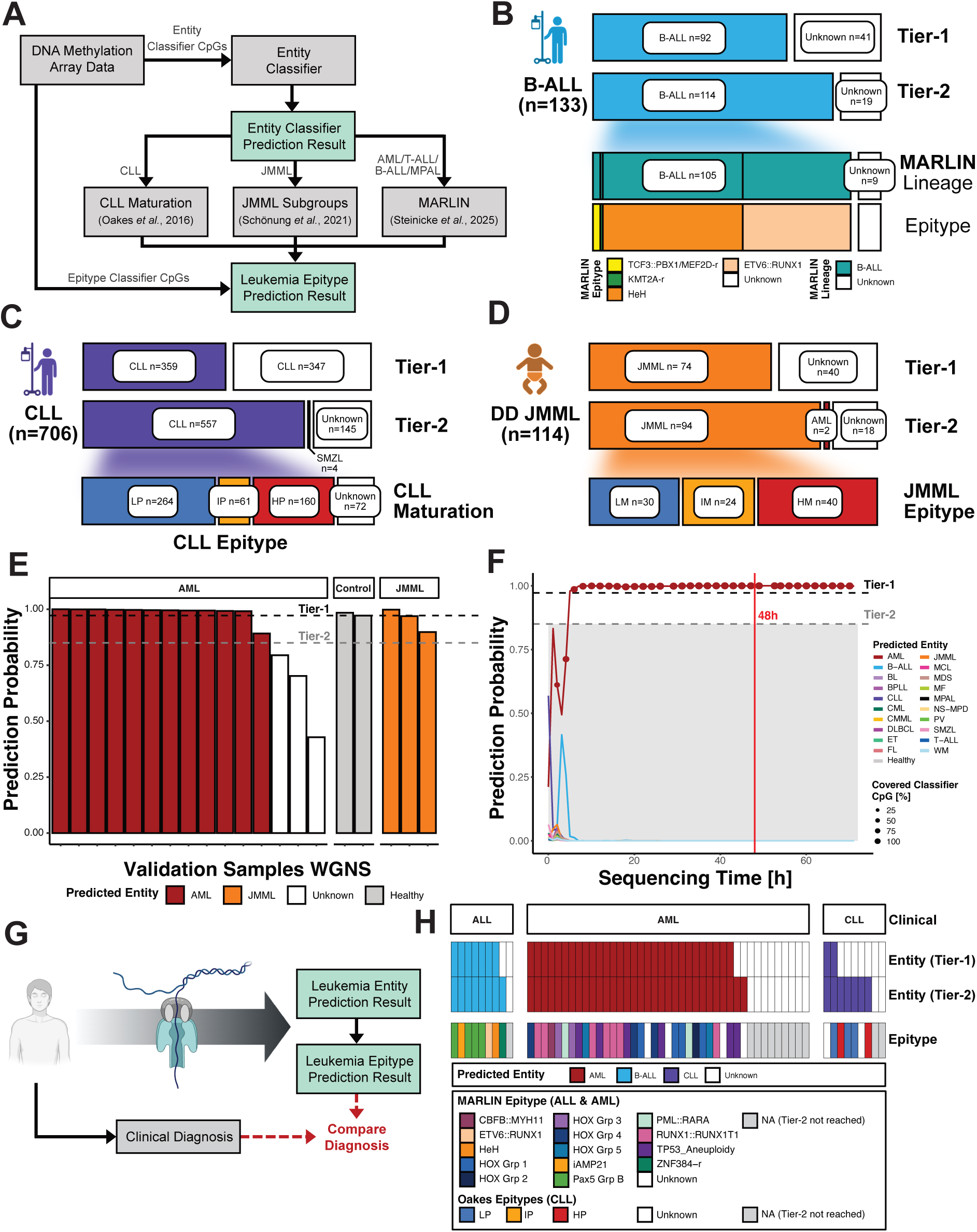
Hierarchical epigenetic classification of disease entity and epitype. (**A**) Schematic overview of the hierarchical classification strategy. **(B-D)** Hierarchical classification was applied to cohorts of 133 B-ALL^26^ (**B**), 706 CLL^27^ (**C**) and 114 patients with a suspected JMML diagnosis submitted to the EWOG-MDS central laboratory (**D**). Entity classification (tier-1 and tier-2 thresholds) results are depicted together with the respective epitype classification for samples reaching a tier-2 threshold. The fraction of samples classified into each disease entity and epitype is shown as bar plots. (**E**) Cross-platform functionality of the entity classifier is shown for a cohort of patients for which we had methylation array and whole genome nanopore sequencing (WGNS) data available (technical validation cohort). WGNS data of AML (n=15), control (n=2) and JMML (n=3) patients was used for classification. Prediction probabilities for the respective entities are plotted. (**F**) Prediction probabilities are plotted at different timepoints of WGNS for a representative AML sample from the technical validation cohort. Colored curves show prediction probabilities for each of the entities included in the classifier and dot sizes represent the cumulative percentage of classifier CpGs covered as a function of sequencing time. (**G**) Hierarchical epigenetic classification was performed in an independent validation cohort consisting of 59 patients. (**H**) Entity and epitype classes were stratified by clinically diagnosed disease entity. MARLIN epitypes were predicted for acute leukemias (ALL and AML) and Oakes epitypes for CLL. Epitype predictions for samples not reaching the tier-2 entity prediction cutoff are assigned as NA.

For CLL, we analyzed data from 706 patients of whom 359 (50.9%) were classified as CLL using the tier-1 threshold and 557 (78.9%) using the tier-2 cutoff (**Figure 3C; Supp. Table 7**). Additionally, 4 patients were classified as SMZL (0.6%) and 145 patients (20.5%) as “unknown” based on the tier-2 cutoff. CLL epitypes were determined using an XGBoost model trained to assign epitypes as previously published by Oakes *et al*.^16^. Of the 557 patients predicted as CLL using the tier-2 threshold, 264 (47.4%) were classified as low-programmed (LP), 61 (11%) as intermediate-programmed (IP) and 160 (28.7%) as high-programmed (HP). A clear epitype assignment was not possible for 72 (12.9%) of the 557 patients classified as CLL, as they did not reach the respective epitype prediction threshold.

Additionally, we analyzed DNA methylation data of 114 patients from the EWOG-MDS study group for which a JMML diagnosis was suspected (**Figure 3D; Supp. Table 8**). Using the tier-1 threshold, 74 patients (64.9%) were classified as JMML. Based on tier-2 threshold, 94 patients (82.5%) received a JMML diagnosis, while 2 patients (1.8%) were diagnosed with AML and 18 patients (15.8%) classified as “unknown”. JMML epitypes^21^ were determined for all patients who received a “JMML” diagnosis. Thirty patients (31.9%) were classified as low methylation (LM), 24 patients (25.5%) as intermediate methylation (IM) and 40 patients (42.6%) as high methylation (HM) JMML. We subsequently collected clinical data for the two patients classified as AML to re-evaluate the suspected diagnosis. The first patient was classified as AML with an entity-level probability of 0.96. The analyzed sample of this patient was collected at JMML relapse after hematopoietic stem cell transplantation (HSCT), at which 18% and 14% bone marrow and peripheral blood blasts were detected, respectively. Molecular analysis revealed a *PTPN11* c.218C>T (p.Thr73Ile) mutation, loss of chromosome 18 and del(6p). Furthermore, analysis of CNVs from DNA methylation array data confirmed del(6p), with additional detection of trisomy 13 and trisomy 21 (**Supp. Figure 3B**). Thus, our entity-level classification together with cytogenetic data hints towards disease progression for this patient.

The second patient was predicted to have AML with a probability of 0.87. This patient had a diagnosis of neurofibromatosis type 1, previously received chemotherapy for astrocytoma, and presented with 50% blasts. Molecular analysis revealed mutations in *PTPN11* c.1508G>C (p.Gly503Ala), *PTPN11* c.1507G>A (p.Gly503Arg), *PTPN11* c.1508G>A (p.Gly503Glu) and *ASXL1* c.2568C>A. Additionally, the patient was found to have a *MECOM*-rearrangement (89% of cells affected), del(17q) and monosomy 7 (**Supp. Figure 3C**). Hence, this patient was accordingly re-classified as therapy-related AML in line with our entity-level classification.

### Hierarchical epigenetic classification using whole genome nanopore sequencing data

DNA methylation classifiers have recently been combined with nanopore sequencing for joint epigenetic and genetic work-ups that can potentially be performed in rapid diagnostic settings^23,28,29^. Hence, we aimed to assess cross-platform functionality of our hierarchical classification strategy using WGNS data as this would enable comprehensive diagnostic molecular profiling in a single assay through the combined reporting of epigenetic entity and epitype classifications together with genetic aberrations. We therefore first established a technical validation cohort by generating WGNS data of 20 patients who have been previously analyzed using DNA methylation arrays. This cohort included 15 AML patients from the ASTRAL-1 trial^30,31^, 3 JMML patients from the EWOG-MDS cohort (array data of one JMML patient was included in classifier training) and two normal controls. To account for the use of this approach on potential low coverage samples and sparse WGNS data, we used a modified entity classifier that was trained on binarized DNA methylation calls. This classifier successfully predicted 17/20 (85%) samples as the correct disease entity using the tier-2 threshold (**Figure 3E**). The remaining 3 (15%) samples did not reach tier-2 threshold and were hence classified as “unknown”. Furthermore, analysis of WGNS data at various sequencing timepoints showed that correct entity-level diagnoses could be established within hours of sequencing, qualifying the classifier for use in a real-time diagnostic setting (**Figure 3F**).

After having established that our entity-level classifier can be applied to WGNS data, we aimed to assess whether our proposed hierarchical classification strategy could recapitulate clinical diagnoses. We therefore analyzed WGNS blinded data from 59 leukemia patients (AML, ALL and CLL) with varying levels of sequencing coverage and compared our DNA methylation-based entity and epitype classifications with clinical diagnoses (**Figure 3G; Supp. Table 9**). The mean WGNS coverage was 10.3 (min: 2.5; max: 19.6) for AML, 2.7 (min: 1.9; max: 3.3) for ALL and 2.9 (min: 2.2; max: 4.4) for CLL (**Supp. Figure 4A**). The mean fraction of covered classifier CpGs were 0.99 (min: 0.87; max: 1.0) for AML, 0.87 (min: 0.74; max: 0.95) for ALL and 0.90 (min: 0.83; max: 0.98) for CLL. A total of 39/59 (66,1%) and 47/59 (79.7%) of the patients were classified above tier-1 and -2 thresholds, respectively (**Figure 3H**). The correct disease entity was predicted for all patients who met the tier-2 threshold (8/9 ALL; 32/41 AML; 7/9 CLL) without misclassifications. There was no significant correlation between entity class prediction probability and sequencing coverage (Pearson’ correlation 0.11; p-value = 0.40) or the fraction of covered classifier CpGs (Pearson’ correlation 0.18; p-value = 0.17). Subsequent epitype predictions were confirmed by clinical diagnostics for 9/10 (90%) of patients assigned to gene fusion-characterized epitypes *RUNX1*::*RUNX1T1*, *PML*::*RARA* and *CBFB*::*MYH11* (**Supp. Figure 4B**). Furthermore, all patients assigned to the *TP53*/Aneuploidy epitype had complex karyotype (CK) AML and all AML patients with *NPM1* mutations were classified as HOXA/B-activated AML epitype which is known be enriched in *NPM1* mutations. Thus, combined DNA methylation-based entity and epitype classification generated results that were compatible with clinically established diagnoses and can be retrieved within hours of nanopore sequencing.

**Figure 4.**
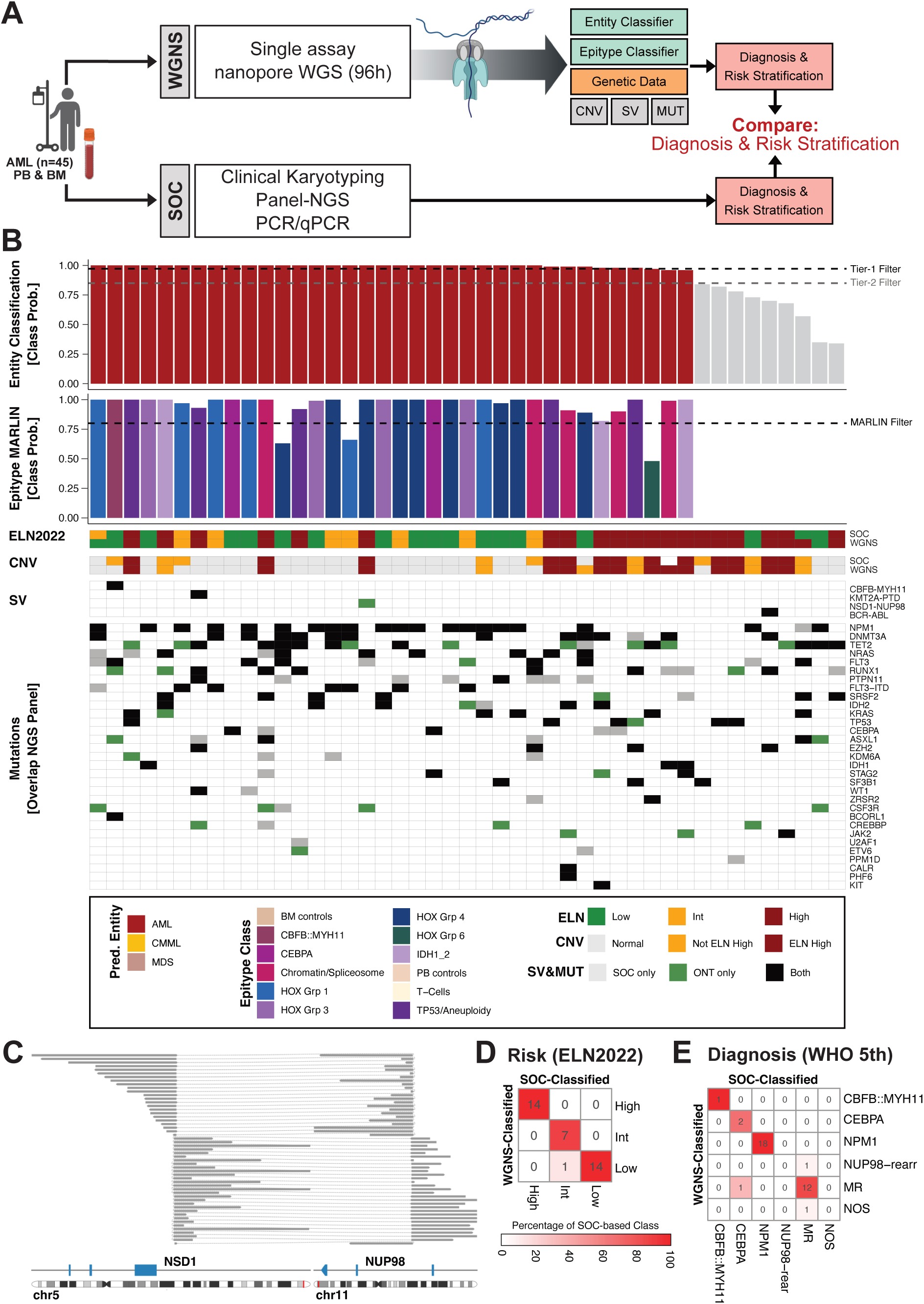
Benchmarking of epigenetic and genetic disease classification against standard-of-care. (**A**) Schematic overview of the benchmarking strategy comparing whole genome nanopore sequencing (WGNS) to standard-of-care (SOC) diagnostics. (**B**) Combined visualization of the epigenetic and genetic diagnosis results. Bar plots showing prediction probabilities for entity and epitype (MARLIN) predictions from WGNS DNA methylation data. Respective probability cutoffs are highlighted as dashed lines. Risk classification (ELN2022), copy number variations (CNVs), structural variants (SVs) and mutations (overlap with panel NGS) are compared between WGNS and SOC. (**C**) Genome browser view of the WGNS detected *NUP98::NSD1* fusion. Supporting split reads are colored in grey and aligned regions of each read connected by dashed lines. (**D-E**) Confusion matrix for comparison of WGNS against SOC based risk stratification (ELN2022) and assignment into diagnostic groups (WHO 5^th^). Heatmap colors indicate the percentage of patients from the SOC-based class.

### Benchmarking of WGNS-based combined genetic and epigenetic diagnostics against standard of care in AML

WGNS has the advantage of analyzing DNA methylation marks and genetic aberrations such as mutations, structural variants (SVs) and copy number variants (CNVs), simultaneously. Analysis of these genetic aberrations is currently crucial for clinical diagnostics and risk stratification. Therefore, we aimed to incorporate this layer into our hierarchical classification framework and to benchmark whether a combination of epigenetic classification together with genetic data from WGNS provides similar diagnostic assignments and risk stratification as the clinical standard of care (SOC). We therefore analyzed either bone marrow or peripheral blood from 45 AML patients using WGNS and compared the obtained results to SOC diagnostics.

Our new diagnostic workflow comprised a single laboratory assay, namely WGNS with a sequencing time of 96h which resulted in a median genome coverage of 35x (min: 12.19; max: 52.77; **Supp. Figure 5A**). This data was used as input for our hierarchical classification framework that relied on DNA methylation-based entity and epitype classification with integration of genetic data. The diagnostic routine for SOC comprised panel sequencing for recurrent AML driver mutations, karyotyping and targeted analysis of gene fusions using PCR and qPCR (**Figure 4A**).

Using our entity classifier, 33/45 (73.3%) and 36/45 (80%) of patients were classified above tier-1 and tier-2 threshold, respectively. All 36 patients who met the tier-2 threshold were classified as AML with no mispredictions (**Figure 4B; Supp. Table 10**). Of the nine patients who did not reach tier-2 accuracy, six out of nine (66.7%) had the highest prediction probability for AML, two for MDS (22.2%) and one for CMML (11.1%) (**Supp. Figure 5B**). Consistent with this, the two patients with the highest class probability for MDS had *TP53* mutations and complex karyotype, while the patient with a high CMML class probability had mutations with VAF ≥ 10% in *ASXL1*, *TET2*, *SRSF2* and *KRAS*, and mutations with VAF < 10% in *FLT3* and *NPM1*. Therefore, the mutational profiles of these three patients are most consistent with the methylation-based entity predictions of MDS and CMML, respectively.

Analysis of MARLIN epitypes revealed assignment of HOXA/B-activated AML for 17 patients, 16 of whom (94.1%) carried an *NPM1* mutation (**Supp. Table 11&12**). The patient without an *NPM1* mutation was assigned to HOX Group 4, which is enriched for *NUP98*-rearranged AML. Consistent with this finding, we detected a *NUP98*::*NSD1* fusion using WGNS that was initially missed by SOC work-up but subsequently could be confirmed at the RNA level using reverse transcription (RT) PCR (**Figure 4C**; **Supp. Figure 6A-D**). Furthermore, all five patients assigned to the *TP53*/Aneuploidy-enriched epitype were diagnosed with myelodysplasia-related AML (AML-MR). SOC work-up detected *TP53* mutations in 2/5 patients (40%), and WGNS in 3/5 patients (60%). Additionally, two of these patients (40%) showed a complex karyotype and one patient had a *KMT2A* partial tandem duplication (PTD) that was detected by both SOC and WGNS workups.

Five patients were classified as Chromatin/Spliceosome-enriched and three as *IDH*-enriched epitypes. For the latter, WGNS and SOC detected mutations in either *IDH1* or *IDH2* in all cases. Additionally, two patients were assigned to the *CEBPA* and one patient to the *CBFB*::*MYH11* epitype. For these three patients, the respective epitype-defining mutations were also detected by both WGNS and SOC. Of note, MARLIN reported a prediction above the published cutoff for all nine patients that did not reach the tier-2 threshold of our entity classifier (**Supp. Figure 5B**). However, three out of nine (33.3%) of these patients were predicted incorrectly as healthy controls (T-cells, BM and PB controls), indicating that our hierarchical classification strategy reduces the number of misclassifications for established epitype classifiers.

Next, we conducted a more detailed comparison of the genomic profiles obtained from WGNS and SOC. WGNS detected a median of 83.3% (min: 40%; max: 100%) of the SNVs identified by SOC work-up per patient whereby VAFs showed a significant correlation across methods (Pearson’s correlation 0.74; p < 2.2×10^−16^; **Supp. Figure 6E**). The majority of SNVs that were missed by WGNS had a VAF of <10% in SOC (**Supp. Figure 6F**). WGNS detected a median of 9 (min: 3; max: 14) additional SNVs per patient within leukemia relevant genes that overlapped with common oncology databases, underlining the advantage of a more comprehensive profiling of mutational landscapes by this technology. Finally, we classified patients according to the WHO definition of AML with defining genetic abnormalities as well as ELN2022 risk groups and compared results to SOC-based assignments (**Figure 4D&E**). Overall, 33/36 (91.7%; Fisher’s test p-value = 1.35 x 10^−12^) patients were assigned to identical WHO groups and 35/36 (97.2%; Fisher’s test p-value = 2.47 x 10^−14^) to identical ELN2022 risk groups. Thus, the epigenetic and genomic classification using WGNS performed comparable to that of SOC.

## DISCUSSION

We here describe the development and validation of a novel hierarchical classification strategy that combines epigenetic and genetic information for molecular work-up in hematologic malignancies. Our strategy explores a new approach to epigenetic diagnostics by first establishing an entity level diagnosis which is then followed by epitype classification choosing appropriate downstream classifiers. This hierarchy of molecular testing enables the use of a single diagnostic platform on which the pretest probability for using a certain epitype classifier is established within the same framework and which does not rely on additional laboratory assays such as flow cytometry or cytopathology. The backbone of this framework is the currently largest collection of DNA methylation data of hematologic malignancies, which we curated as a reference atlas to train our entity-level classifier. This collection encompasses 21 diseased and healthy states and comprises a comprehensive resource of hematopoietic neoplasm data sets. As we needed to rely on the, sometimes sparsely, annotated patients’ diagnoses in publicly available datasets, these annotations might not necessarily represent the perfect ground truth and probably hinder the development of a classifier with a 100% accuracy. In addition, diagnostic criteria, e.g. for AML and MDS, have changed over the last years which might further impact on the precision in differentiating these entities using the available databases. We tried to address this issue by training our classifier model on large patient numbers to prevent overfitting on individuals with misdiagnoses. By validating our classifier in independent patient cohorts, we were able to prove the success of our strategy by high diagnostic accuracies and the ability of our classifier to identify patients with likely inaccurate diagnoses.

We established two prediction cut-offs (tier-1 and -2) for an entity-level diagnosis. Although these cut-offs reduce the number of patients that can be successfully assigned to disease entities, they provide an important quality measure to ensure that predictions reach clinically acceptable accuracy, specificity, and sensitivity values. We currently recommend to follow tier-1 predictions and use additional clinical and genetic data for patients where the tier-2 cut-off but not tier-1 is fulfilled. Prediction probabilities that are below the tier-2 cut-off can be explained by several scenarios: i) samples have a low tumor-cell content which might cause an intermediate methylation profile between tumor and normal; ii) the full heterogeneity of disease entities which are only represented by a low number of patients in our training dataset might not be covered which results in low prediction probabilities; iii) patients are not treatment naïve which might impact on classifier performance; iv) patients with overlap syndromes such as MPN/MDS overlap disorders^32^; v) disease entities which are not currently included in our classifier training. It will be a task for future work to generate additional large datasets for entities which are underrepresented or not represented at all in our classifier to further improve the performance of our diagnostic classifier model.

We validated the functionality of our classification strategy across different technological platforms, namely DNA methylation arrays and nanopore-sequencing. In combination with our classifier, the latter technology allows for rapid diagnostic work-ups as an entity level diagnosis can already be established within hours of sequencing. A similar strategy has recently been clinically validated for rapid work-up of acute leukemias and CNS tumors, including intra-operative nanopore-sequencing for surgical decision making^23,28^. Importantly, combining DNA methylation classifiers with WGNS enables combined epigenetic and genetic work-up. The latter is currently required for establishment of an accurate diagnosis for many hematologic malignancies according to WHO definitions and key for identification of suitable targeted therapies and risk group assignment^2,3,5^. Here, we benchmarked for the first time this combined work-up for hematologic neoplasms in a cohort of AML patients and proved non-inferiority of our single platform hierarchical WGNS testing strategy in assigning WHO diagnoses and ELN2022 risk groups compared to SOC. Additionally, we identified a diagnosis-defining *NUP98*-rearrangement that was missed by SOC work-up, thus underscoring the power of long-read WGNS for detection of SVs. Yet, a limitation of our WGNS approach was the low sensitivity for the detection of SNVs with low VAFs which is likely caused by differences in sequencing coverage between panel-sequencing and WGNS. However, this limitation might be overcome by future studies that combine whole genome read-outs with adaptive sampling or CRIPSR/Cas9 enrichments for an improved coverage at genomic loci with recurrent mutations in hematologic malignancies.

The integrative profiling of genetic and epigenetic markers will likely gain additionally significance in the next decades as an increasing number of molecular alterations are discovered that serve as biomarkers for targeted therapies^33^. Rapid profiling of these alterations will become a central part of the initial diagnostic work-up and will likely be subject of future clinical trials. We demonstrate that our newly developed hierarchical testing strategy can serve as a versatile framework for single platform diagnostic work-up which can be easily amended with additional epitype classifiers that might be developed in the future.

## METHODS

### Data collection and patient samples

DNA methylation array data of 47 datasets from patients with hematopoietic neoplasm and healthy controls were collected (**Supplementary Table 1**). Data from diagnostic timepoints were selected for datasets where this information was annotated. As independent validation cohorts, we obtained DNA methylation array data for 133 B-ALL (GSE56600) and 706 CLL cases^27^. Additionally, DNA methylation array data was generated from bone marrow (BM) or peripheral blood (PB) DNA of 114 patients with a suspected JMML diagnosis from the EWOG-MDS study group. Illumina Infinium HumanMethylation450 (450k) and MethylationEPIC (EPICv1 and v2) BeadChip array data were generated at the Molecular Profiling and Screening Core Facility of the German Cancer Research Center (DKFZ). WGNS was performed on samples from the ASTRAL-1 study, the EWOG-MDS study group, Charité - Universitätsmedizin Berlin (Department of Hematology, Oncology and Cancer Immunology), the DKFZ/NCT/DKTK MASTER program (NCT05852522), and the Heidelberg Cell and Liquid Biobank (Department of Hematology, Oncology and Rheumatology) of Heidelberg University Hospital^30,31,34,35^. Informed consent of patients or legal guardians was obtained and the study conducted in accordance with the Declaration of Helsinki. Generation, collection and storage of patient data was approved by institutional ethics committees.

### Quality control and processing of DNA methylation array data

Quality control of DNA methylation array data was conducted using adapted versions of *conumee2* (https://github.com/mwsill/conumee2) and *minfi* (https://github.com/mwsill/minfi) that were optimized for cross platform DNA methylation array analysis including EPICv2^36,37^. First, the average of the log2 median of methylated and unmethylated channels per sample was calculated and samples with a value of <10.5 removed. Second, we calculated the array noise per sample using the *conummee2* “CNV.fit” function and removed samples with a value of >0.5. DNA methylation array processing was performed using *minfi* (https://github.com/mwsill/minfi)^36^. We first applied a probe-level filtering and removed probes with detection p-values ≥0.01, masked probes across any DNA methylation array version, probes mapping to sex chromosomes or common SNPs and non-CpG targeting probes. Probe annotations, masking and filtering criteria were derived from https://zwdzwd.github.io/InfiniumAnnotation^25^. We furthermore included only CpG sites that were covered across all Illumina DNA methylation array version (450k, EPICv1 and EPICv2). DNA methylation beta values were calculated from probe intensity levels and probes with “NA” values in any sample removed. DNA methylation array data was normalized using beta-mixture quantile (BMIQ) normalization as implemented in the *watermelon* (version 2.12.0) R package^38,39^. As patient samples might be duplicated in publicly available datasets, we additionally applied a correlation filter to remove closely identical DNA methylation arrays. For this, we calculated sample-wise correlation values across the 5000 most variable CpG probes based on standard deviation. Samples were considered duplicated if Pearson’s correlation values exceeded 0.999, and only one of the affected samples was retained in the dataset.

### Dimensional reduction analysis

For dimensional reduction analysis, CpG sites with a standard deviation of more than 0.25 across all samples were selected (n=12,627). Principal component analysis (PCA) was calculated and the number of PCs that contribute to a cumulative proportion of variance of 0.8 selected (n=34). Uniform Manifold Approximation and Projection (UMAP) was calculated based on the selected PCs using the R *umap* (version 0.2.10) package with default parameters changes for n_neighbours=30 and min_dist=0.5^40^. UMAP coordinates were plotted using *ggplot2* (version 3.5.2)^41^.

### Differential methylation and ChromHMM enrichment analysis

Differential methylation analysis was conducted to determine differentially methylated probes (DMPs) between i) healthy cell types and ii) hematopoietic neoplasms using the *minfi* (https://github.com/mwsill/minfi) “dmpFinder” function^36^. DMPs were subsequently selected based on an absolute mean beta value difference of >0.2 between groups and q-value of <0.05. The overlap between both sets was visualized using the *UpSetR* (version 1.4.0) package^42^. Enrichment for ChromHMM states was conducted based on publicly available probe-level annotations https://zwdzwd.github.io/InfiniumAnnotation. Each DMP set was considered as foreground (FG) while probes that remained after quality filtering in our DNA methylation array dataset but did not overlap with DMPs were considered as background (BG). The log2 enrichment fold-change of proportion of probes in FG overlapping with each ChromHMM state compared to BG was calculated and significance determined using Fisher’s exact test. Results were plotted using *ggplot2*^41^.

### DNA methylation classifier training

For classifier training, we focused on the previously determined 12,627 CpG sites with a standard deviation of more than 0.25 across all samples. Data from healthy cell types were removed from the dataset. The data was then split into a training (n=3,884) and testing (n=958) cohort using the *caret* (version 7.0-1) “createDataPartition” function with a training fraction of 0.8^43^. An Extreme Gradient Boosting Model (XGBoost) was trained for multi-class classification using the *caret* “xgbTree” method^44^. Hyperparameter tuning was conducted using a 5-fold cross validations strategy followed by model training using the following settings: nround=270, eta=0.2, max_depth=1, gamma=0.5, colsample_bytree=0.4, min_child_weight=2 and subsample=0.5. Feature selection was performed by applying the *caret* “varImp” function and CpG sites with a feature importance of >0.1 selected (n=932). A final model was trained based on the previously determined hyperparameters and selected CpG sites. ROC curves were calculated using the *pROC* (version 1.19.0.1) R package based on binarized predictions (concordance with annotated diagnosis) and maximum class probabilities to identify the “tier-1” threshold where the classifier reports with optimal specificity and sensitivity^45^. The “tier-2” threshold was defined based on a tradeoff analysis between valid patient predictions and classifier accuracy. Alluvial plots were generated using the *ggalluvial* (version 0.12.5) R package and heatmaps using *pheatmap* (version 1.0.12)^46,47^. For cross-platform functionality, data was binarized using a beta-value cutoff of 0.6 and an XGBoost model trained based on all 12,627 varCpGs with default hyperparameters (nround=100, eta=0.3, max_depth=6, gamma=0, colsample_bytree=1, min_child_weight=1 and subsample=1) to prevent overfitting of a classifier on CpG subsets when analyzing sparse data.

### Library preparation and WGNS of samples from Charité Berlin

DNA was extracted from patient samples of a retrospective cohort (comprising CLL, B-ALL and AML) using an AllPrep DNA/RNA kit (QIAGEN), following the manufacturer’s instructions. DNA quality was analyzed using TapeStation (Agilent Technologies). Libraries were prepared using Ligation Sequencing kits SQK-LSK109 or SQK-LSK110 for sequencing on MinION R9.4.1 flow cells or SQK-LSK114 for sequencing on PromethION R 10.4.1 flow cells (Oxford Nanopore Technologies; **Supp. Table 9**). Library preparation was thereby performed following the manufacturer’s instructions using 1100 ng of input DNA and Long Fragment Buffer (LFB). Libraries were sequenced for 24h.

### Library preparation and WGNS of AML samples for benchmarking

Quality control of DNA was conducted using TapeStation (Agilent). If necessary, DNA was sheared to fragments in a range of 20 to 25 kb using g-TUBEs (Covaris) as described previously^48^ and DNA fragments below 10 kb were removed by size selection using the Short Read Eliminator (SRE) XS kit (Pacific Biosciences). Nanopore sequencing libraries were prepared using the Ligation Sequencing Kit V14 (SQK-LSK114, Oxford Nanopore Technologies) according to manufacturer’s instructions and sequenced on a PromethION 24 (Oxford Nanopore Technologies) device using FLO-PRO114M flow cells for 96h. Nuclease washes using the Flow Cell Wash Kit EXP-WSH004 (Oxford Nanopore Technologies) were carried out when necessary, according to the manufacturer’s instructions, followed by the resumption of sequencing from recovered or fresh library material.

### WGNS data analysis

Alignment of WGNS data against the hg38 reference genome was conducted using *dorado* (version 1.0.2; https://github.com/nanoporetech/dorado), with a min-qscore filter of 10 and the “dna_r10.4.1_e8.2_400bps_hac@v5.0.0” and “dna_r10.4.1_e8.2_400bps_hac@v5.0.0_5mCG_5hmCG@v3” models for base and modified base calling. Samples sequenced on R9.4.1 flow cells were processed using “dna_r9.4.1_e8_hac@v3.3” and “dna_r9.4.1_e8_hac@3.3_5mCG” models. Bam files were sorted and indexed using *samtools* (version 1.20)^49^. For analysis of DNA methylation data *modkit* (version 0.4.3; https://github.com/nanoporetech/modkit) “pileup” was used for construction of bedMethyl files, with the “--cpg”, “--combine-strands” and “--combine-mods” flags, and a target bed file specifying genomic coordinates of CpG sites that are included on the 450k array. Data was subsequently binarized and transformed into input formats for entity-level or epitype classifiers. Tumor-only SNV calling was performed using ClairS-TO (version 0.4.2) with min_coverage of 3, indel_min_af of 0.05 and “ont_r10_dorado_sup_5khz” platform specification^50^. Initial variant filtering was conducted using BCFtools (version 1.20) requiring a quality score cutoff of more than 10, sequencing depth of more than 10 reads, and a minimum of 3 supporting variant reads with a variant allele frequency of at least 0.05^49^. Variant annotation was performed using vcf2maf (https://github.com/mskcc/vcf2maf), Ensembl VEP, and OncoKB annotator (https://github.com/oncokb/oncokb-annotator)^51,52^. Variants with an allele frequency of <0.05 in both the “1000 Genomes Project” and gnomAD that were either included in COSMIC or OncoKB and overlapped with commonly mutated genes in hematopoietic malignancies (**Supp. Table 13**) were reported ^52–56^. Additionally, variants detected by SOC work-up which were not included in the automated pipeline reports were manually inspected for presence in WGNS data.

Tumor-only SVs were determined based on a recently developed consensus strategy (https://www.biorxiv.org/content/10.1101/2025.07.30.667694v1.full). Therefore, SVs were called using both *delly* (version 1.5.0) in longread (lr) mode and *Severus* (version 1.5) with the provided hg38 variable number tandem repeats (VNTR) and 1000 Genomes Project panel of normal (PON) files^57–59^. The *sansa* (version 0.2.5;https://github.com/dellytools/sansa) “compvcf” function (flags --nosvt -e 0 -m 0 -n 250000000) was subsequently used to identify SVs that were detected by both tools followed by filtering for somatic only SVs. Likely remaining germline SVs with a VAF of 1.0 were excluded and SVs filtered for variants with more than 3 supporting reads. SVs overlapping with a previously curated candidate list or recurrent variants in AML were reported^5,60^.

Internal tandem duplications (ITDs) in FLT3 (FLT3-ITD) were furthermore analyzed using *breaktracer* (version 0.0.8; https://github.com/tobiasrausch/breaktracer) with two different settings (-m 5 -c 5 -s 5 -r 0 and -m 15 -c 15 -s 15 -r 0). CNV calling was conducted using QDNAseq as implemented in the „epi2me-labs/wf-human-variation” (https://github.com/epi2me-labs/wf-human-variation) pipeline and by disabling coverage filtering, followed by annotation of CNVs to cytogenetic bands ^61^.

### Epitype classifier

#### JMML

An updated version of the JMML epitype classifier was trained which featured analysis of 450k, EPICv1 and EPICv2 data as the previously published version was not functional for EPICv2 due to differences in the array design. Therefore, the definition of JMML epitypes by Schönung *et al*. was considered as a gold standard and the new classifier version trained based on the published batch corrected data^21^. For feature selection, the mean DNA methylation beta-value for each probe stratified by epitype was calculated and the 2000 CpG sites with the largest absolute mean difference between LM and IM, LM and HM, IM and HM identified (n= 3391 unique CpGs). For classifier development, data was randomly split into a training (n=229) and a testing (n=55) cohort using the *caret* “createDataPartition” function and an XGBoost model trained^43^. Hyperparameter tuning was performed using a 5-fold cross validation strategy, resulting in the best accuracy for nrounds = 30, eta = 0.1, max_depth = 5, gamma = 0, colsample_bytree = 0.8, min_child_weight = 1 and subsample = 0.75. A final JMML classifier model was trained based on CpGs with a feature importance greater than 0.1 (n=121).

#### CLL

A CLL epitype classifier was developed based on a previously published dataset and annotated epitypes^16^. Data was therefore randomly split into a training (n=121) and a testing (n=38) cohort using the *caret* “createDataPartition” function and 1,048 CpGs selected that overlapped with a previously determined set of probes localized at transcription factor (TF) binding sites of six CLL relevant TFs^16,43^. An XGBoost classifier was trained based on these sites with default hyperparameters (nround=100, eta=0.3, max_depth=6, gamma=0, colsample_bytree=1, min_child_weight=1 and subsample=1) using the *caret* “xgbTree” method^44^. ROC curves were calculated using the *pROC* R package based on binarized predictions (concordance with annotated epitype) and maximum class probabilities to identify a suitable threshold for epitype classification with high specificity and sensitivity^45^. Thereby, a CLL epitype threshold of 0.78 was determined. For analysis of WGNS data, the classifier was adjusted by training based on binarized DNA methylation data with a beta-value cutoff 0.6.

#### MARLIN-RF

The MARLIN Random Forest (RF) classifier was developed in R (v4.4.1), following a similar approach to Capper *et al.*^7^ and using the acute leukemia dataset established by Steinicke *et al*.^23^. Input features were chosen from the union of the 5,000 most variable CpGs from each of (i) the full acute leukemia reference cohort, (ii) AML methylation classes (excluding *PML::RARA*, *RUNX1::RUNX1T1* and *CBFB::MYH11*) together with BM and PB controls, (ii) B-ALL methylation classes together with B-cell controls and (iv) T-ALL methylation classes together with T-cell controls, yielding 14,700 input features in total. For final model training, an RF with 10,000 trees was trained on the full reference cohort using the *randomForest* (version v4.7-1.1) R package^62^, with per-tree downsampling to the smallest class to adjust for class imbalance. In a second step, raw RF scores were recalibrated to class probabilities by fitting an L2-penalized multinomial logistic regression model using the *glmnet* (version v4.1-8) R package^63^, trained on independent RF scores obtained from a 5-fold cross-validation. To estimate classifier performance, nested 5×5-fold cross-validation was used, with RF training and score recalibration performed exclusively on the training partition of each outer fold. The inner folds were used to generate independent RF scores for fitting the calibration model, which was then applied to the corresponding outer-fold test predictions. Similar to the MARLIN neural network version, biologically related classes were aggregated into methylation class families and a prediction threshold of 0.8 was used. For analysis of WGNS data, MARLIN-based epitype classification was performed as previously described using the neural network version^23^.

### RT-PCR validation NUP98-rearrangement

RNA of patients was reverse transcribed to cDNA using the “High Capacity cDNA Reverse Transcription Kit” (Applied Biosystems) according to the manufacturer’s instructions. PCR amplification of fusion products or control amplicons was performed using AmpliTaq Gold (Applied Biosystems) with established primers for NUP98::NSD1 fusions and NUP98 controls (**Supp. Table 14**)^64^. PCR conditions were adjusted to 95°C for 10 min, followed by 40 cycles of 95°C for 45 sec, 62°C for 45 sec and 72°C for 1 min, followed by 72°C for 10min. PCR products were analyzed on an agarose gel and by Sanger sequencing.

### Data and code availability

DNA methylation array data of public datasets were gathered as specified in **Supp. Table 1**. DNA methylation array and WGNS data generated in the course of this study were uploaded to “The German Human Genome-Phenome Archive” (GHGA; https://www.ghga.de/) under accession number **XYZ** (will be provided upon publication). Bioinformatic analysis scripts are available on GitHub (https://github.com/MaxSchoenung/nanoleukemia).

## AUTHOR CONTRIBUTIONS

M.S. and D.B.L. designed and coordinated the study. M.S., M.T., P. Lajer, A.D., J.H., K.R., L.K., C.R. performed experiments. M.S., C.G., C.P., L.B. and D.B.L. supervised experiments and/or provided materials and reagents. M.S., T.R., T.L.S., E. Sträng, M.S.O. and L.K. performed bioinformatic analysis and/or developed software. S.R., E. Sollier, T.S., J.C.S., M.D-F., C.C.O., J.I.M-S., U.G., M.D., A.C., C.F., C.M.N., H.D., F.H.H., C.M.-T., M. Hundemer, M. E., L.B. provided and/or collected patient samples and/or data. M.S., P.H., V.H., S.F., R.F.S., J.K., M. Hartmann, P. Lutsik, M. Hundemer, M.E., L.B., and D.B.L. interpreted the data and/or supervised analysis. M.S. and D.B.L. wrote the initial draft of the manuscript. All authors contributed and agreed with the final version of the manuscript.

## ACKNOWLEDGEMENTS

We thank the German Cancer Research Center (DKFZ) Molecular Profiling and Screening Core Facility, Omics IT and Data Management Core Facility (ODCF), NGS Core Facility and NCT Sample Processing Laboratory for excellent technical support. We furthermore thank the EWOG-MDS study group for supporting this project. Figure subpanels were created in BioRender: Hartmann, M. (2026) https://BioRender.com/i91831i.

## SUPPLEMENTARY FIGURES

**Supplementary Figure 1.**
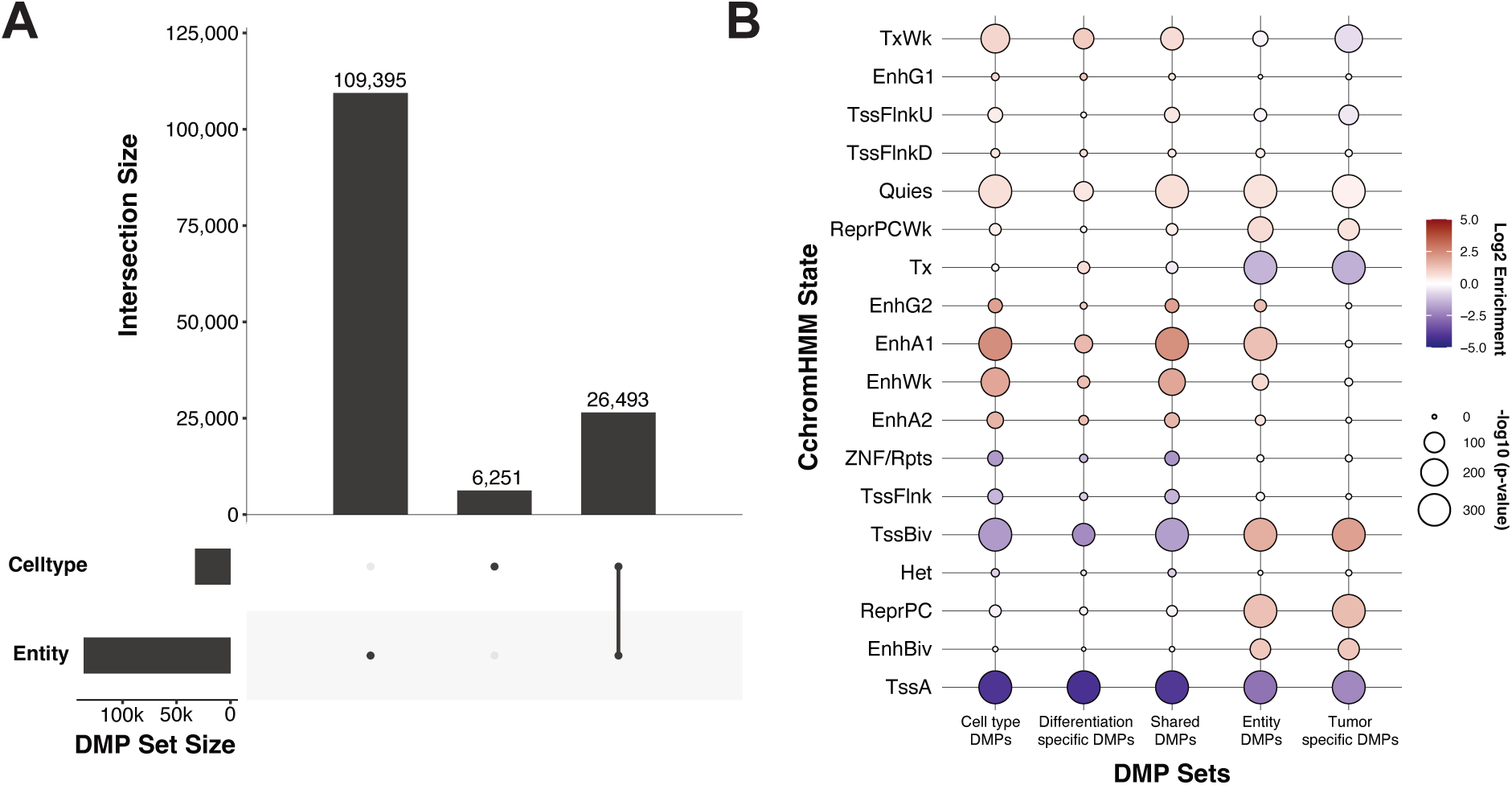
Differentiation- and entity-specific DNA methylation programs regulate distinct genomic regions. **A)** Upset plot visualizing the overlap between cell type and entity DMPs. **B)** Enrichment analysis of ChromHMM states for cell type and entity DMPs, as well as unique (differentiation and cell type-specific) and shared DMPs between both sets. Colors show log2 enrichment values over the background of included probes in the dataset and the circle sizes the negative log10 p-value of Fisher’s test.

**Supplementary Figure 2.**
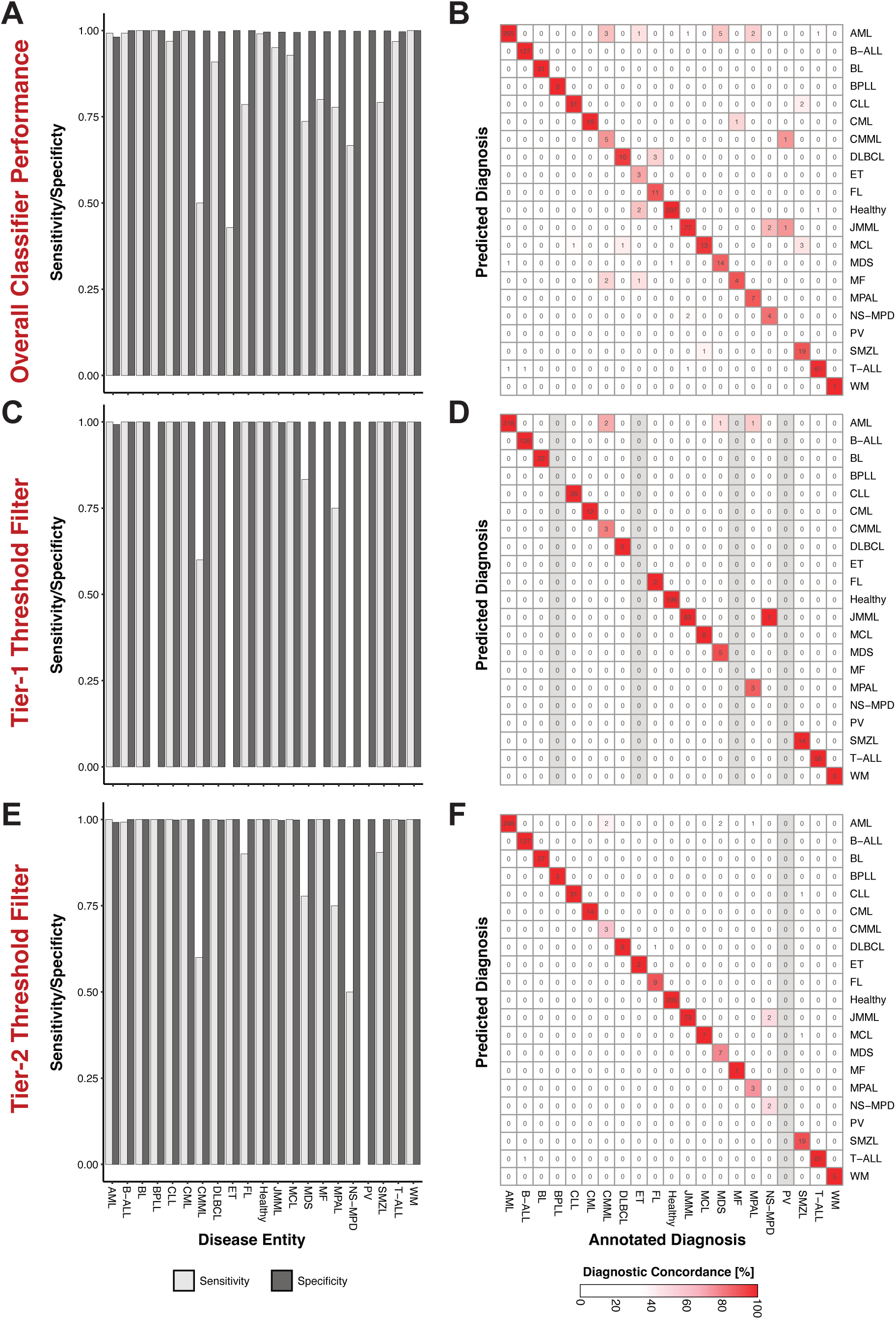
Overview of classifier performance. Different classifier cut-offs were analyzed using per class sensitivity and specificity (**A, C, E**) and confusion matrices (**B, D, F**). The overall classifier performance without filtering (**A&B**) was thereby compared to the tier-1 (**C&D**) and tier-2 (**E&F**) thresholds. The color of the confusion matrix indicates the diagnostic concordance as percentage of the annotated diagnosis and grey boxes classes where patients have been excluded by filtering.

**Supplementary Figure 3.**
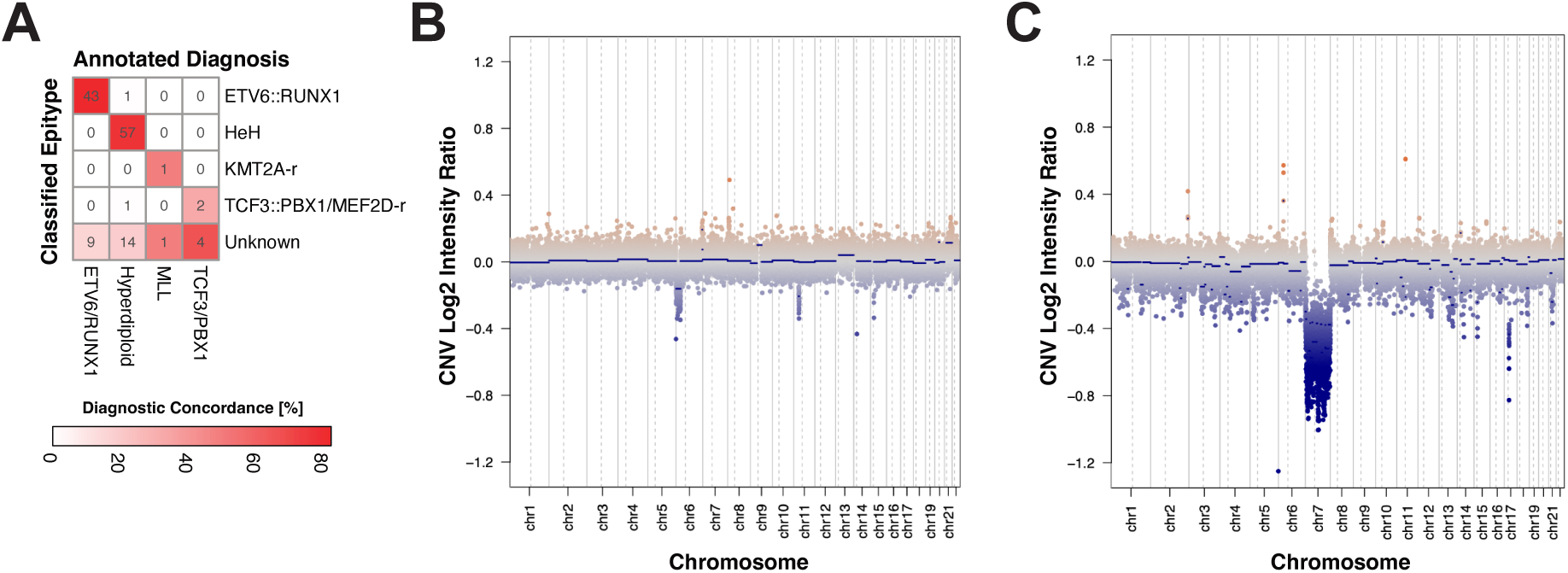
Genetic alterations in the validation cohorts. **A)** Confusion matrix comparing the annotated genetic alterations to classified epitypes. **B-C**) CNV plot for EWOG-MDS patients that were classified as AML.

**Supplementary Figure 4.**
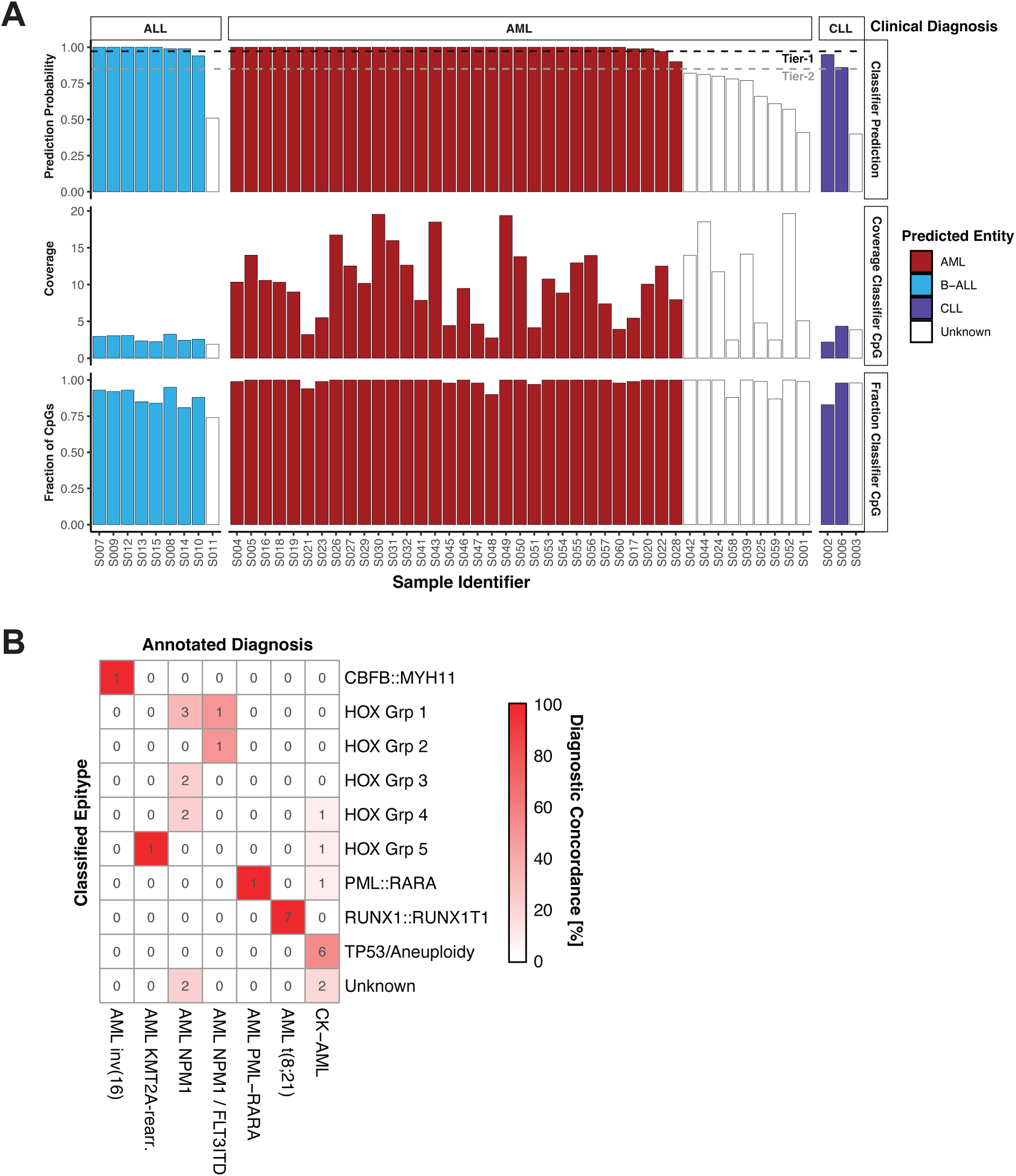
Classification of WGNS data from leukemia patients. **A)** Classifier probabilities and sequencing statistics for the 59 patients analyzed by WGNS stratified by clinical diagnosis. Class probability for the entity with the highest score is shown together with sequencing coverage at classifier CpGs and the fraction of classifier CpGs covered by each WGNS run. Tier-1 and tier-2 thresholds are annotated. **B**) Confusion matrix comparing the annotated genetic alterations to classified epitypes for patients that were predicted as AML using the tier-2 threshold.

**Supplementary Figure 5.**
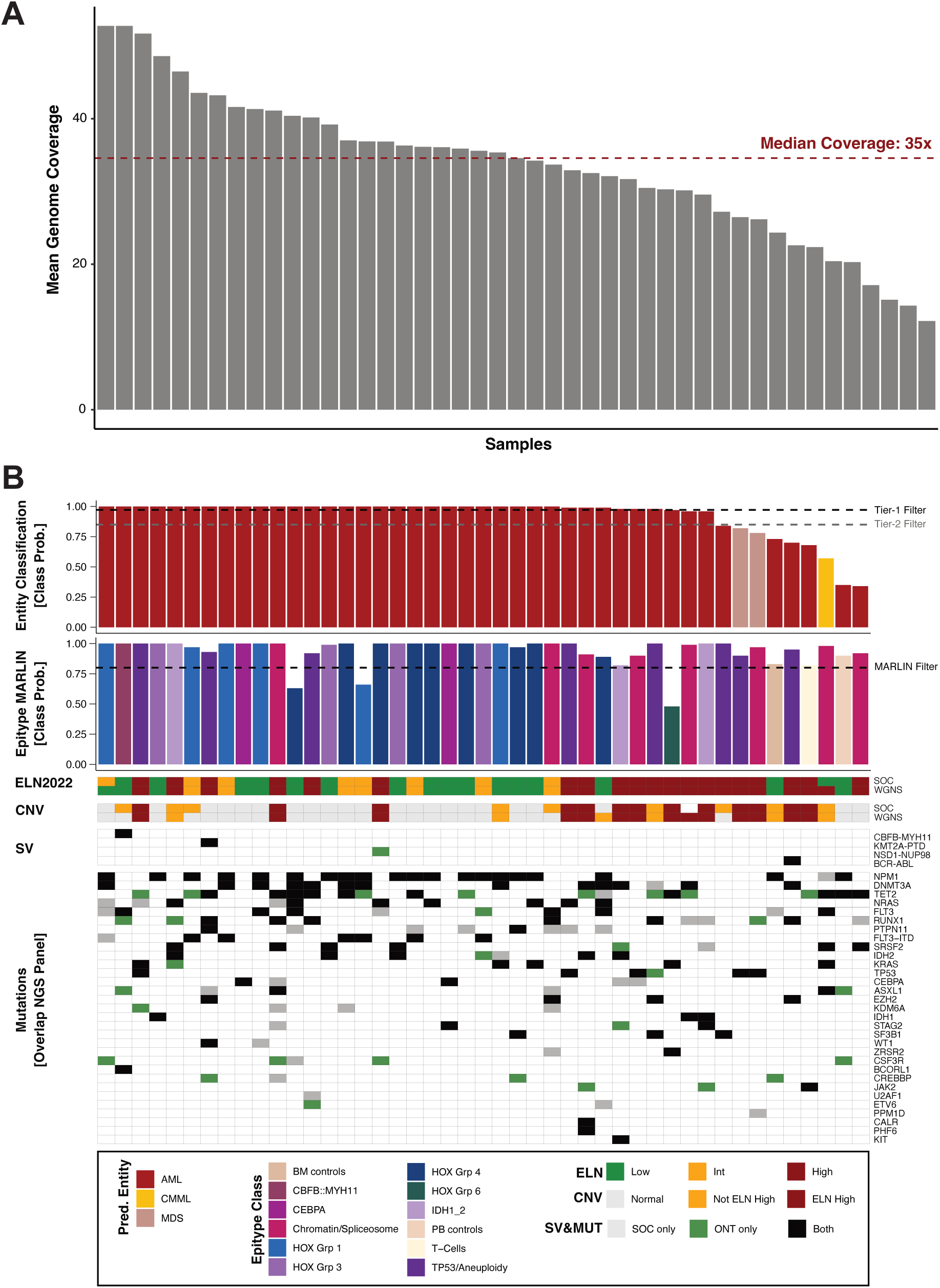
Sequencing coverage and predictions for AML WGNS patients. **A)** Mean genome coverage of WGNS patients included in the study. The median coverage across all samples is highlighted. **B**) Combined visualization of the epigenetic and genetic diagnosis results. Bar plots showing prediction probabilities for entity and epitype (MARLIN) predictions from WGNS DNA methylation data. Respective probability cutoffs are highlighted as dashed lines. Risk classification (ELN2022), copy number variations (CNVs), structural variants (SVs) and mutations (overlap with panel NGS) are compared between WGNS and SOC.

**Supplementary Figure 6.**
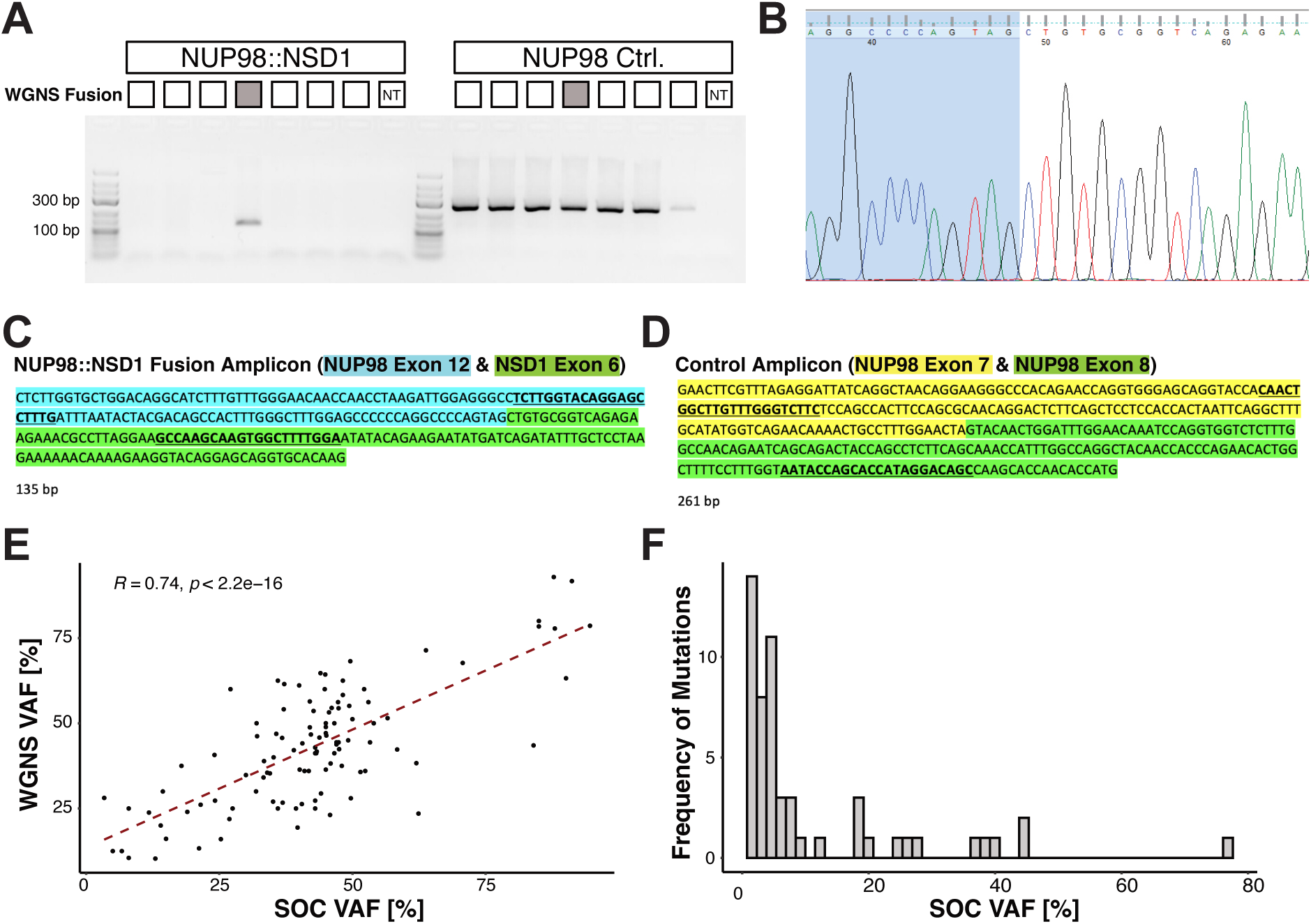
Detection of genomic alterations by WGNS. **A)** RT-PCR for detection of the NUP98::NSD1 fusion. Agarose gel for the RT-PCR products (NUP98::NSD1 fusion detection and NUP98 control) for 6 WGNS patients and a non-targeting (NT) control. The patient with the NUP98::NSD1 fusion detected by WGNS is highlighted. **B**) Sanger sequencing track confirming the translocation between NUP98 exon 12 and NSD1 exon 6. **C-D**) Predicted amplicon sequences for the (**C**) fusion product and (**D**) control amplicon. (**E**) Pairwise comparison and correlation between VAFs calculated by WGNS and SOC. Pearson’s correlation and correlation test p-value are annotated. (**F**) Histogram showing the SOC VAF for mutations that are not detected by WGNS.

